# Environmentally enriched pigs have transcriptional profiles consistent with neuroprotective effects and reduced microglial activity

**DOI:** 10.1101/262469

**Authors:** SM Brown, SJ Bush, KM Summers, DA Hume, AB Lawrence

## Abstract

Environmental enrichment (EE) is widely used to study the effects of external factors on brain development, function and health in rodent models, but very little is known of the effects of EE on the brain in a large animal model such as the pig. Twenty-four young pigs (aged 5 weeks at start of study, 1:1 male: female ratio) were housed in environmentally enriched (EE) pens and provided with additional enrichment stimulation (a bag filled with straw) once daily. Litter, weight and sex matched controls n= (24) were housed in barren (B) conditions. Behaviour was recorded on alternate days from study day 10. After 21 days, RNA-sequencing of the frontal cortex of male piglets culled one hour after the enrichment stimulation, but not those at 4 hours after stimulation, showed upregulation of genes involved in neuronal activity and synaptic plasticity in the EE compared to the B condition. This result is mirrored in the behavioural response to the stimulation which showed a peak in activity around the 1 hour time-point. By contrast, EE piglets displayed a signature consistent with a relative decrease in microglial activity compared to those in the B condition. These results confirm those from rodents, suggesting that EE may also confer neuronal health benefits in large mammal models, through a potential relative reduction in neuroinflammatory process and increase in neuroprotection driven by an enrichment-induced increase in behavioural activity.

## 1. Introduction

Environmental enrichment (EE) paradigms are widely used in laboratory rodents as a method to study the effects of environmental influences on brain development and function (1). EE utilises multiple methods of environmental modification to facilitate cognitive and motor stimulation in a species appropriate manner. In rodents this is often achieved through the use of novel objects, running wheels, nesting material and group housing (2).

The beneficial effects of EE on physiology and brain function in rodents are well documented (reviewed in (3)). EE paradigms have been shown to have a neuroprotective effect through the inhibition of spontaneous apoptosis (4) and through increased cell proliferation and neurogenesis in the hippocampus (5–7) and prefrontal cortex (8). Indeed, EE has been shown to limit neurodegeneration in models of aging (9, 10), and can rescue some behavioural and physiological brain effects of early life stress (11, 12). Current evidence suggests this is most likely precipitated by the mechanisms underlying neuronal plasticity, due to increased expression of trophic factors and immediate early genes (IEG) in the brains of EE animals (13–15).

Expression of neurotrophins has also been proposed as a contributory factor in mood disorders, in particular stress-induced depression and anxiety. While the main focus has been around expression of brain derived neurotrophic factor (BDNF) (16–18), there are a number of other trophic factors and IEGs with a role in stress resilience and mood. Neuronal growth factor (Ngf) mRNA has been shown to be reduced in the dentate gyrus in response to restraint stress in rats (19). Environmental deprivation during early life in rats reduces IEG expression in the prefrontal cortex both as a chronic effect (20), in response to social interaction (21), and after chronic social defeat stress in mice(22). Levels of IEG Arc mRNA in the frontal cortex and hippocampus of rats exposed to predator scent stress were reduced in those animals which displayed an extreme anxiety-like behavioural response, and in the dentate gyrus Arc mRNA expression levels also correlated with levels of circulating corticosterone(23).

Interestingly, depressive-like behaviour in rodents induced by social isolation in early life not only results in decreased expression of a number of IEG involved in synaptic plasticity, including Bdnf, Egr-1 and Arc ((24), mice), but also in increased microglial activation ((25), rats). Microglia are specialised macrophages of the central nervous system that play a role in neuromodulation, phagocytosis and inflammation (reviewed in(26)). There is increasing evidence to suggest that microglial mediated brain inflammation is a contributing factor to a number of mental health and nervous system disorders, such as multiple sclerosis(27), Alzheimer’s disease(28), schizophrenia (29, 30), and major depression(31). While the exact mechanism by which microglia contribute to these disorders is not yet known, psychological stress has been shown to result in increased levels of microglia in the prefrontal cortex of rats (32) while early life stress, in the form of maternal deprivation, has also been shown to lead to increases in microglial motility (33) and phagocytic activity (34) that persist into adulthood. Anti-inflammatory drugs such as minocycline (which blocks microglial activation) produce antidepressant effects in rodent models (35). Several anti-psychotic medications have also been shown to block microglial activation (36, 37).

The physiological and behavioural effects of a number of nervous system disorders can be modulated by EE in rodent models (38), and EE is now widely used as an experimental tool in studies of neurodegeneration and neural development (39). Due to its beneficial effects on behaviour, EE has long been used as a method to improve the welfare of production animals, by providing environmental manipulations that allow the animal to behave in a more species-typical way (reviewed in (40)). However there has been little work on the effects of EE on the brain in a large animal model, such as the pig. The aim of the present study was to investigate the effects of EE on gene expression within the frontal cortex of young pigs with the aim of identifying potential gene expression signatures of a healthy brain in a large animal model, and to potentially identify early signatures of neural ill-health in the absence of abnormal behaviours.

## 2. Materials and Methods

### 2.1 Ethical review

All work was carried out in accordance with the U.K. Animals (Scientific Procedures) Act 1986 under EU Directive 2010/63/EU following ethical approval by SRUC (Scotland’s Rural College) Animal Experiments Committee. All routine animal management procedures were adhered to by trained staff and health issues treated as required. All piglets remaining at the end of the study were returned to commercial stock.

### 2.2 Animals and general experimental procedures

Post-weaning behavioural observations were carried out on litters from six commercial cross-bred mothers (Large White x Landrace); the boar-line was American Hampshire. Artificial insemination was performed using commercially available pooled semen. Litters were born within a 72 hour time window in free-farrowing pens which allowed visual and physical contact (nasal contact through bars) with neighbouring litters. No tooth resection was performed and males were not castrated. In line with EU Council Directive 2008/120/EC tail docking is not routinely performed. Weaning occurred at 24-27 days of age. At weaning piglets were weighed, vaccinated against Porcine Circoviral Disease and ear tagged for identification. Post-wean diet was in the form of pelleted feed (Primary Diets Prime Link Extra) provided ad libitum. At 4 days post weaning, 8 piglets per litter were selected as being as close to the litter average weight as possible while balancing for sex (1:1 male to female). These 8 piglets per litter were split into sex balanced quartets to give final group sizes of 4 piglets per matched group. One group was then housed in an enriched (EE) pen and one in a barren (B) pen, all in the same room (Figure 1). Post weaning pens did not allow for physical or visual contact between neighbours. All pens contained feed troughs ~80cm in length and piglets had access to one water nipple per pen.

**Figure 1:**
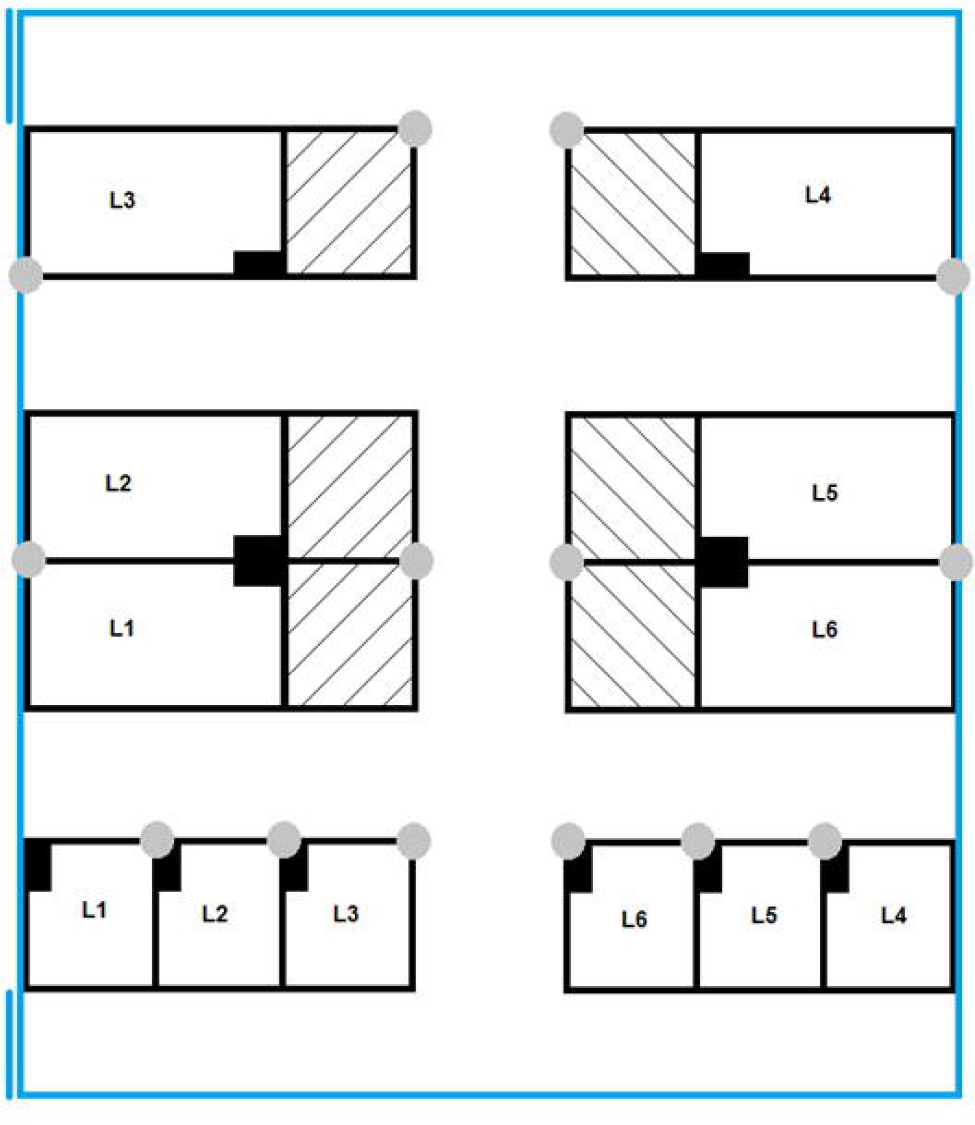
Diagram of the layout of the experimental room. Double lines signify the outer room doors used for access. Grey circles indicate the location of cameras and filled black boxes show the location of feeders. Litter of origin is identified as L1-L6. Empty pens are indicated by hatched lines. EE pens were twice the size of B pens as indicated in the diagram. All pens were solid floored. EE pens had the addition of straw over two thirds of the flooring.

**Figure 2:**
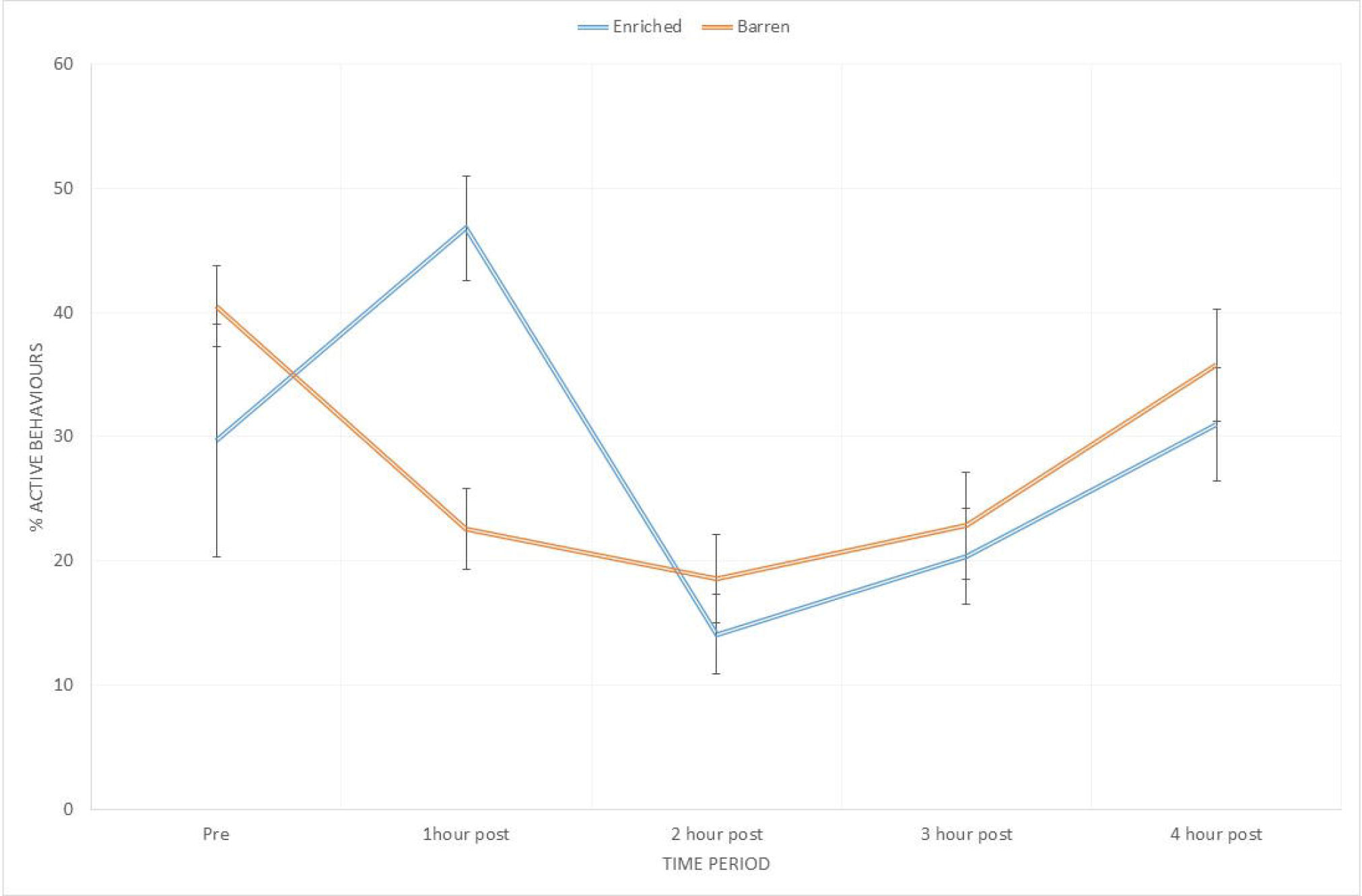
Percentage of behaviours classified as ‘active’ at each time point for EE and B piglets. Values are means with error bars indicating SEM. 60 scans per pen contribute to each time point after the addition of the bag (at time 0) and 15scans per pen in the pre period. Active behaviours include locomotion (without object or social) (running, walking, hopping, pivoting and flopping), object interaction (with enrichment, i.e. bedding or enrichment stimulus, or components of pen) and nosing (of pen-mates or pen structure). X-axis shows the time relative to the provision of the additional enrichment stimulus.

All piglets were marked on the back with spray marker paint to allow individual identification. B pens measured 1.8m x 1.8m with concrete floors and rubber matting (1.8m x 1.2m) in one corner to provide some thermal insulation. Kennels were provided for the first 10 days to enhance thermal comfort as B housed piglets had no opportunity to use straw to control their thermal environment as the EE pigs could. EE pens measured 3.6m x 1.8m also with concrete flooring and with large quantities of straw covering the floor (covering half the pen to a depth of 0.15m). The same rubber matting was also provided in the enriched pens. Piglets in the six EE pens were provided with additional enrichment once daily (enrichment stimulus) in the form of a plastic bag (approximately 400mm x 750mm x 150mm) filled with straw (approximately 2kgs). This bag was placed in the EE pens after morning husbandry was complete (at around 10 am). At the same time the gate of the matched B pen was approached to account for the stimulating effect of human presence. The bag was removed at the end of each day (~1700 hours) and a new bag used on the following day. The bag was open at one end to allow straw to be extracted by the piglets and was the only source of fresh straw given to the EE pigs daily. The bag was first introduced at day 5 post weaning and thereafter daily. Pen cleaning and enrichment order was randomised daily. In accordance with the Defra Code of Recommendations for the Welfare of Livestock(41), temperature within the room was automatically controlled at 22°C and artificial lighting was maintained between the hours of 0800 to 1600, with low level night lighting at other times.

### 2.3 Behavioural observations

Piglets were digitally recorded in their home pen using Sony LL20 low light cameras with infra-red and a Geovision GV-DVR. Two cameras were set up per EE pen, one at the rear and one at the front to provide maximal coverage, and one camera per B pen. Piglets were observed in their home pens on five days between the ages of 41 and 53 days (+/-3 days due to staggered farrowing times). Piglets were all observed on the same days during the same time periods to negate any time of day effects on sampling. The behaviour scores are the cumulative total of how many times that behaviour occurred in the timeframe within the pen. Observations began after morning husbandry was complete, 15 minutes prior to the time point when the additional enrichment was provided to the EE pigs, and finished 4 hours after the provision of additional enrichment. Behaviours were recorded by scan sampling with a frequency of one minute(42) for the total 4.25hour duration. The observer used a 10 second window to aid in accurate identification of the behaviour occurring at the start of every scan sample. A condensed ethogram based on previous work in enrichment (43) and play (44) in pigs (Table 1) was used. All behaviours were recorded as frequencies. One observer completed all video analysis to remove any reliability issues relating to multiple observers.

**Table 1:** The ethogram used for behavioural scoring. Individual postures were considered inactive behaviours and all other behaviours as active as when active the posture is included in the active behaviour description and not scored separately.

| *Category/Behaviour* | *Definition/type* |
| --- | --- |
| **Active Behaviours** |  |
| **Social** |  |
| Belly nosing penmate | A distinctive, rhythmic up-and-down movement of a piglet's snout against the belly of another piglet. Piglets may be standing, sitting or lying when performing or receiving belly nosing |
| Tail biting | Manipulation, sucking or chewing tail of pen mate |
| **Locomotion (without object or social)** |  |
| Walking | A steady 4 beat gait with 2 or 3 legs bearing weight at any one time (depending on the phase of the movement). |
| Running | Energetic running and hopping in forward motions often associated with excitability, using large areas of the pen, occasionally coming into marginal/accidental contact with other piglets |
| Hop/flop/pivot | High energy movements on the spot in an upwards (hop), downwards (flop) or circular (pivot) direction but with little or no forward propulsion |
| **Object and pen Interaction** |  |
| Object interaction_enrichment stimulus | Any active interaction with the enrichment stimulus. Piglet may be locomotory or stationary. Includes biting/chewing of bag, walking or running with bag, climbing in bag, shaking bag vigorously and lying on bag. |
| Object interaction_no enrichment stimulus | Active interaction with other pen components, such as straw, rubber matting and shavings. As above may include walking or running with the object, and chewing/biting object |
| Nosing/exploring pen | Piglet walking/running or stationary with nose to floor or pen walls performing a gentle nosing motion |
| Feeding/drinking | Piglet oriented with snout in trough/drinker appearing to be chewing or swallowing |
| ***Inactive Behaviours*** |  |
| **Postures** |  |
| Lateral lying | Piglet lying on its side |
| Ventral lying | Piglet lying on its front |
| Sitting | Piglet sitting on its haunches with front legs straight |
| Standing | Piglet standing still. Not performing any other visible behaviour |
| **Other** |  |
| Not visible | Piglet out of view of camera |

### 2.4 Statistical analysis of behaviour

The full dataset was condensed to give one data point for each behaviour per hour period per day for each pen being the cumulative recordings for all piglets in that pen in that observation period. The introduction of the bag is time-point 0. Analysis was performed on these mean values for each period per pen averaged across the 5 observation days. Analysis of the behavioural data was performed in Minitab version 17.3 (2016) using a generalized linear model with treatment as a fixed effect and time point as a random effect. Sex, litter of origin and pen were included in the model as covariates for the behaviour (and weight data).

### 2.5 Tissue collection

At 54 days of age (+/- 2 days) one male piglet per pen was sedated at one hour after the introduction of the additional enrichment stimulus, and one male per pen at 4 hours post enrichment stimulus. An intramuscular injection was used with a combination of Domitor (1mg/ml medetomidine hydrochloride) (0.01ml/kg), Ketamine (0.1ml/kg), Hypnovel (5mg/ml medazolam) (0.1ml/kg) and Stresnil (40mg/ml azaperone) (0.025ml/kg) (as previously described in 43). Piglets were left for 15 min to allow sedation to take full effect before being euthanized via intercardial injection with Euthatal (200mg/ml sodium pentobarbital) (0.7ml/kg) for brain tissue collection. Male brain tissue was sampled to be consistent with other EE studies, mainly performed in rodents. Piglet brains were removed whole and dissected over dry ice. For continuity all dissections were performed by a single researcher using http://www.anatomie-amsterdam.nl/sub_sites/pig_brain_atlas for reference, utilising both parasagittal and rostrocaudal views. Frontal cortex was placed in RNAlater (Ambion) and agitated on a tube rotator for 15 minutes to assist in the perfusion of RNAlater through the tissue. Samples were frozen at −20˚C until required.

### 2.6 Preparation of tissue samples

Frontal cortex samples in RNAlater were removed from −20˚C and allowed to reach room temperature. RNA extraction was performed using the Qiagen RNeasy lipid tissue kit as per manufacturer’s instructions. 80mg of tissue was used from each sample and a TissueRupter (Qiagen) used for homogenization. Extracted RNA was stored at −20˚C until required (<3months). Samples were quantified and quality checked using a 2200 Tapestation (Agilent).

### 2.7 RNA-seq and differential expression analysis

Poly A+ library preparation and RNA sequencing (RNA-seq) of the prepared samples was carried out by Edinburgh Genomics Next Generation Sequencing Service (http://genomics.ed.ac.uk/services/sequencing) using the Illumina HiSeq 4000 platform, generating >30 million strand-specific 75bp paired-end reads per sample. RNA-seq reads were processed using the high-speed transcription quantification tool Kallisto(45), version 0.42.4, generating transcript-level expression estimates both as transcripts per million (TPM) and estimated read count. Kallisto quantifies expression by building an index of k-mers from a set of reference transcripts and mapping the reads to these directly. For this purpose, we used the combined set of cDNAs and ncRNAs from annotation Sscrofa11.1 (ftp://ftp.ensembl.org/pub/release-90/fasta/sus_scrofa/cdna/Sus_scrofa.Sscrofa11.1.cdna.all.fa.gz and ftp://ftp.ensembl.org/pub/release-90/fasta/sus_scrofa/ncrna/Sus_scrofa.Sscrofa11.1.ncrna.fa.gz, respectively, accessed 1^st^ November 2017). Transcript-level read counts were then summarised to the gene level using the R/Bioconductor package tximport v1.0.3 (46), with associated HGNC gene names obtained from Ensembl BioMart v90 (47). The tximport package collates Kallisto’s output into a count matrix, and calculates an offset to correct for differential transcript lengths between samples (the consequence of differential isoform usage). The count matrix was imported into the R/Bioconductor package edgeR v3.14.0 (48) for differential expression analysis, with the ‘trimmed mean of M values’ method used to normalise gene counts. Genes with expression level estimates lower than 2 TPM in one or both treatment groups were filtered out as noise. After filtering lowly expressed genes, the final dataset comprised 21,971 genes. Genes were considered to be differentially expressed (EE vs B) when they returned a between group fold-change greater than +/-1.2, with a *p*-value<0.02. These thresholds for fold change and p-value were chosen specifically for this study, to minimise noise with regard to identifying a conservative set of significantly differentially expressed genes. A fold change of +/- 1.2 reduces the likelihood of natural variation in expression (i.e. which would be observed around a fold change of 1), as does an alpha level more stringent than the conventional 0.05. The sequencing data generated for this project is deposited in the European Nucleotide Archive (ENA) under study accession number PRJEB24165 (http://www.ebi.ac.uk/ena/data/view/PRJEB24165).

### 2.8 Gene set and gene signature enrichment analysis

Gene set enrichment analysis was performed on the differentially expressed gene lists using Panther.db v13.0 (release 20170413) with default parameters. To assess the statistical significance of enrichment-induced variation in gene expression for small groups of genes with a common function (‘signature genes’), we used a randomisation test as in (49). Two sets of gene signatures, derived from previous human studies, were first obtained for IEG genes (e.g. (50–58)) and microglial genes (59) (Table S4), and filtered to remove those genes not annotated in pigs (resulting in n = 123 and 569 genes, respectively). Subsets of x genes were drawn at random s = 10,000 times from the set of all genes for which a fold change is quantified (i.e. for each comparison of treatment groups), where x = number of signature genes. We calculated q, the number of times the set of signature genes had a higher (for IEG genes) or lower (for microglial genes) median fold change than each randomly chosen subset. Letting *r = s-q*, then the *p*-value of this test is *r+1/s*+1.

### 2.9 Network analysis of transcriptomic changes

These expression values were entered into the network analysis tool Graphia Professional (formerly Miru, derived from BioLayout Express^3D^ (60, 61); http://kajeka.com/graphia-professional). A sample-to-sample analysis was performed using a correlation coefficient threshold of 0.99 to assess the impact of individual variation on all gene expression profiles. A gene-to-gene network was also created from the whole dataset using a correlation coefficient threshold of 0.90. The nodes (genes) were clustered using the Markov Clustering (MCL) algorithm (62) with an inflation value (which determines cluster granularity) of 2.2. This algorithm identifies tightly coordinated sub-structures within the overall network.

## 3. Results

### 3.1 Additional enrichment stimulation increases active behaviour

An analysis of the percentage of active behaviours in EE vs. B housed piglets found a treatment by time-point effect (F_4,43_=4.2, *p*=0.005). The coefficients show this difference to be attributable to an increased proportion of active behaviours in the EE piglets at one hour post provision of the enrichment stimulus (t_4_=3.99, *p*<0.001)(Figure 2). Piglet sex was not observed to have an effect on the active behavioural response to enrichment.

A housing treatment x time effect was observed in locomotor (without object or social) behaviour (F_4,43_=3.19, *p*=0.021), interaction with the enrichment stimulus (F_4,43_=59.42, *p*<0.001) and interaction with other pen components, such as rubber matting (F_4,43_=2.75, *p*=0.038). These effects peaked at one hour post provision of the enrichment stimulus to the EE pens with greater performance of locomotor (without object or social) behaviour (t_4_=2.75,*p*=0.008) and interaction with the enrichment stimulus (t_4_=15.08, *p*<0.001), and at two hours post enrichment stimulus for interaction with other pen objects (t_4_=2.07, *p*=0.04) in the B pens. A greater proportion of lateral lying in B housed pigs was observed at one hour post enrichment stimulus (t_4_=2.69, *p*=0.01). An effect of housing treatment alone was observed in nosing of pen-mates (F_1,47_=25.9, *p*=0.007) with more nosing of pen-mates occurring in the B housed piglets. An effect of time alone was found in feeding behaviour (F_4,43_=37.13, *p*=0.002), nosing of the pen (F_4,43_=11.47, *p*=0.018) and a trend towards lateral lying (F_4,43_=4.3, *p*=0.09). No effect of time or treatment was observed in any of the other behaviours recorded. Means and S.E.M for behaviours are shown in table S1.

There was no difference between piglet growth (as measured by Average Daily Gain in weight, ADG) during the pre-weaning period, prior to the start of the study (F(1, 47) = 0.027, *p* = 0.873). Piglet growth was observed to differ between environments (F(1, 47) = 5.68, *p* = 0.038) during the study period, with EE piglets having on average a higher ADG than B piglets (EE mean = 688.9g SEM = 15.5, B mean = 654.0g SEM = 12.6).

### 3.2 Transient up- and downregulation of genes in response to environmental enrichment

Complete lists of differentially regulated genes are provided in Table S2. Initial analysis revealed that for a large majority of genes, expression in the frontal cortex was unaffected by the environmental enrichment (Figure 3).

**Figure 3:**
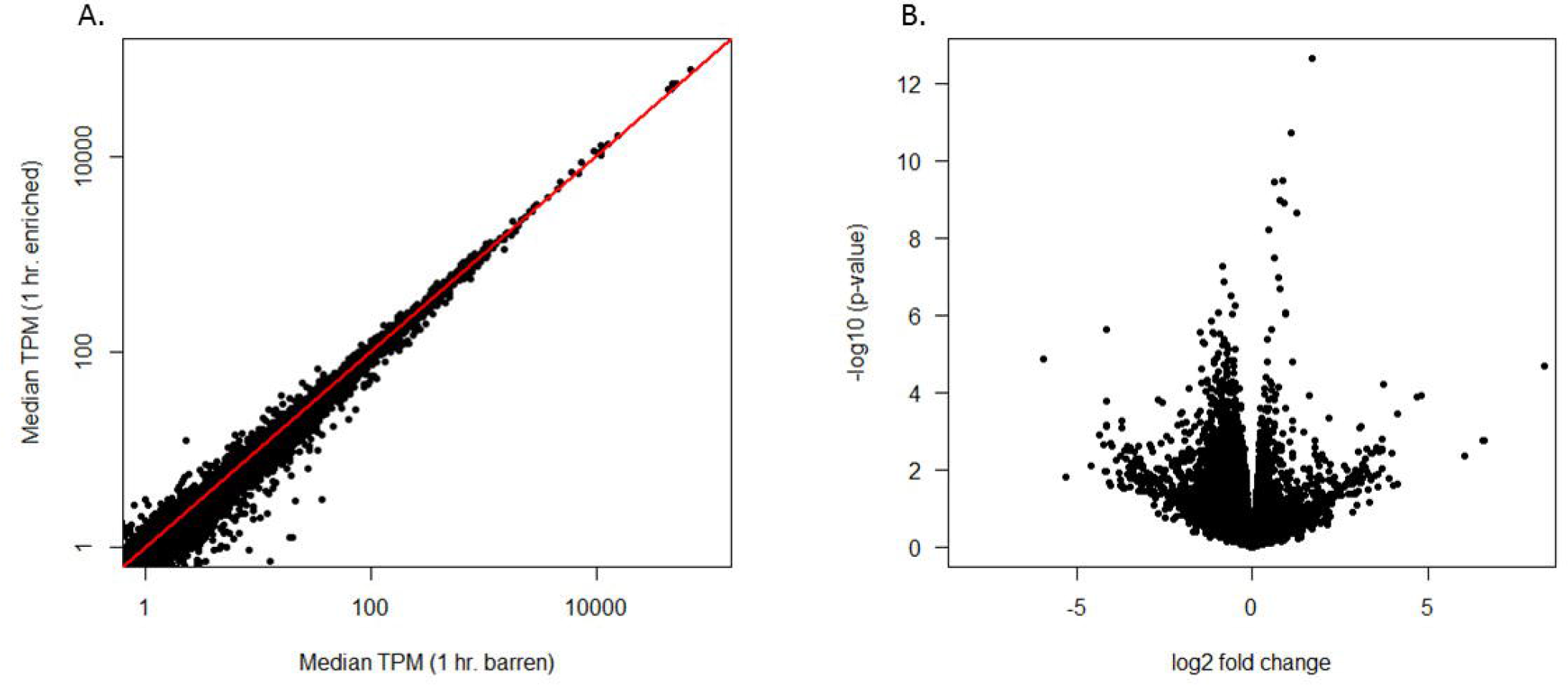
Differentially expressed genes. (Panel A) Largely similar expression profiles of frontal cortex in barren (x-axis) and enriched (y-axis) environments, with the exception of a subset of differentially expressed genes. Each point is a gene, with expression level measured as the median transcripts per million (TPM) across all replicates. The line y = x is shown in red. Differentially expressed genes are more distant from this red line. (Panel B) Volcano plot of the fold change in expression for all genes. X axis shows log2 fold change; Y axis shows –log10 *p*-value. Few genes demonstrate large fold changes, with most clustering at the root of the graph.

When all 21,971 genes were included in a network analysis (see Materials and Methods), members of the same litter showed similar expression patterns, regardless of treatment or time. This was particularly noticeable for litters L3, L4 and L6 (Figure 4A). Using the full gene set there was no association with treatment (Figure 4B) or time (not shown).

**Figure 4:**
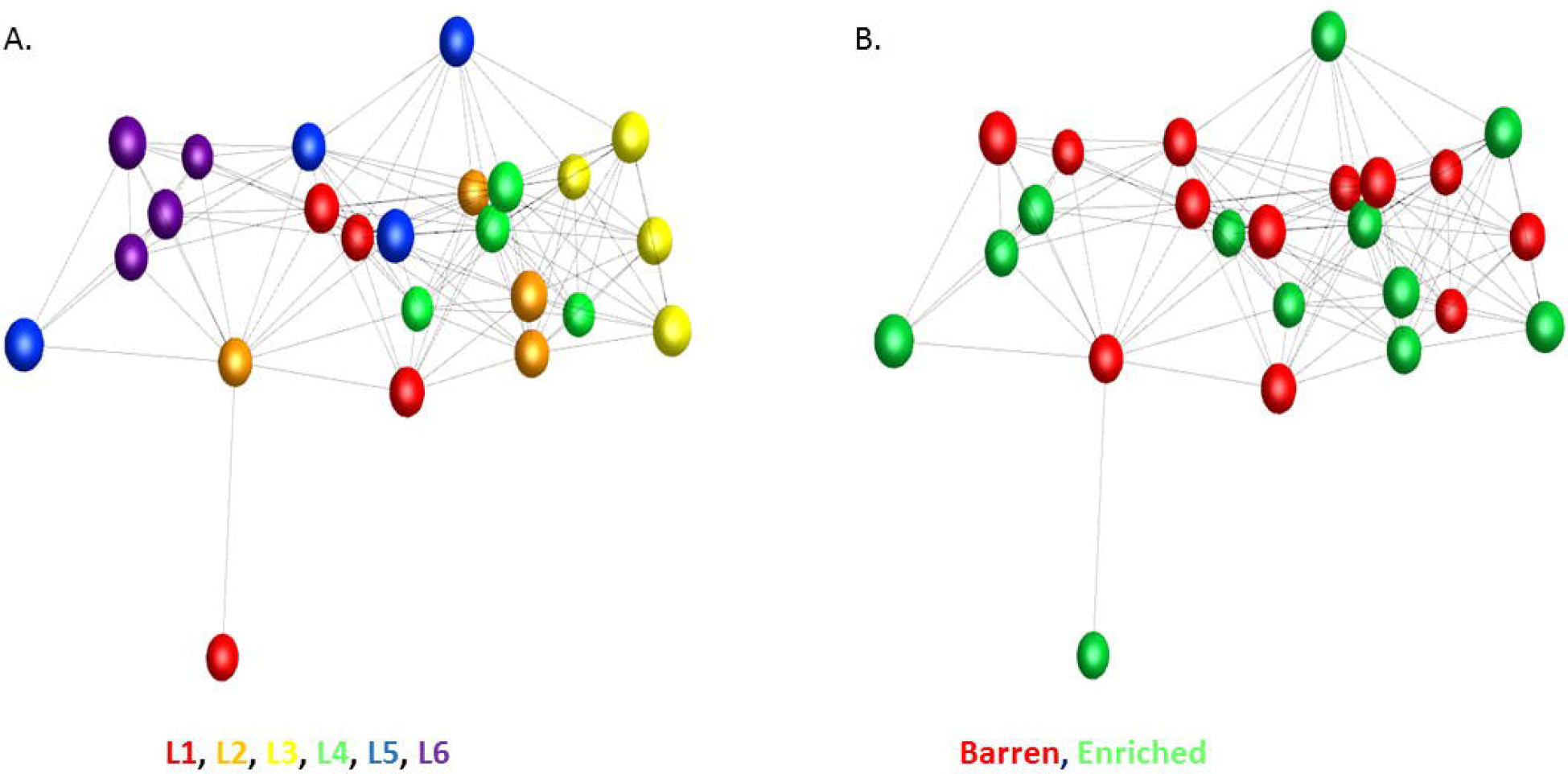
3D network graph of all genes expressed in the frontal cortex. Each node (sphere) in the network graph represents a sample (individual), and each edge (line) a Pearson’s correlation between samples ≥ 0.99. The local structure of the graph shows that gene expression profiles cluster according to litter of origin (Panel A, where nodes are coloured by litter) and that there is little effect of treatment (i.e. no clear clustering) upon overall gene expression (Panel B, where nodes are coloured by environment).

A set of 349 genes were up-regulated after one hour of environmental enrichment but appeared to return to baseline after four hours (Table 2). A second set of genes was down-regulated at both one hour and four hours (Table 2) in the EE cohort. The composition of these two gene sets is discussed below.

**Table 2:** Numbers of genes differentially expressed in the frontal cortex of piglets housed in EE and B environments at one and four hours post enrichment. The last two rows combine all piglets within each treatment regardless of time-point.

| Comparison | Direction of Change | # Differentially Expressed<br>Genes |
| --- | --- | --- |
| 1hr EE vs. 1Hr B | Up | 357 |
|  | Down | 652 |
| 4hr EE vs. 4hr B | Up | 46 |
|  | Down | 83 |
| All EE vs. All B | Up | 125 |
|  | Down | 184 |

Gene set enrichment analysis at one hour post enrichment highlights an over-representation of genes involved in synaptic transmission being up-regulated in the EE animals (Table 3), with an over-representation of genes involved in apoptosis and cellular defence being down-regulated in the EE animals. Further analysis of the gene lists revealed that approximately 8% of those genes upregulated at 1 hour in the EE animals are known IEGs while 15% of those genes down-regulated in the EE (up-regulated in B) animals at 1 hour are related to microglial processes. At four hours post enrichment no gene sets were found to be overrepresented using the Panther GSEA overrepresentation test.

**Table 3:**
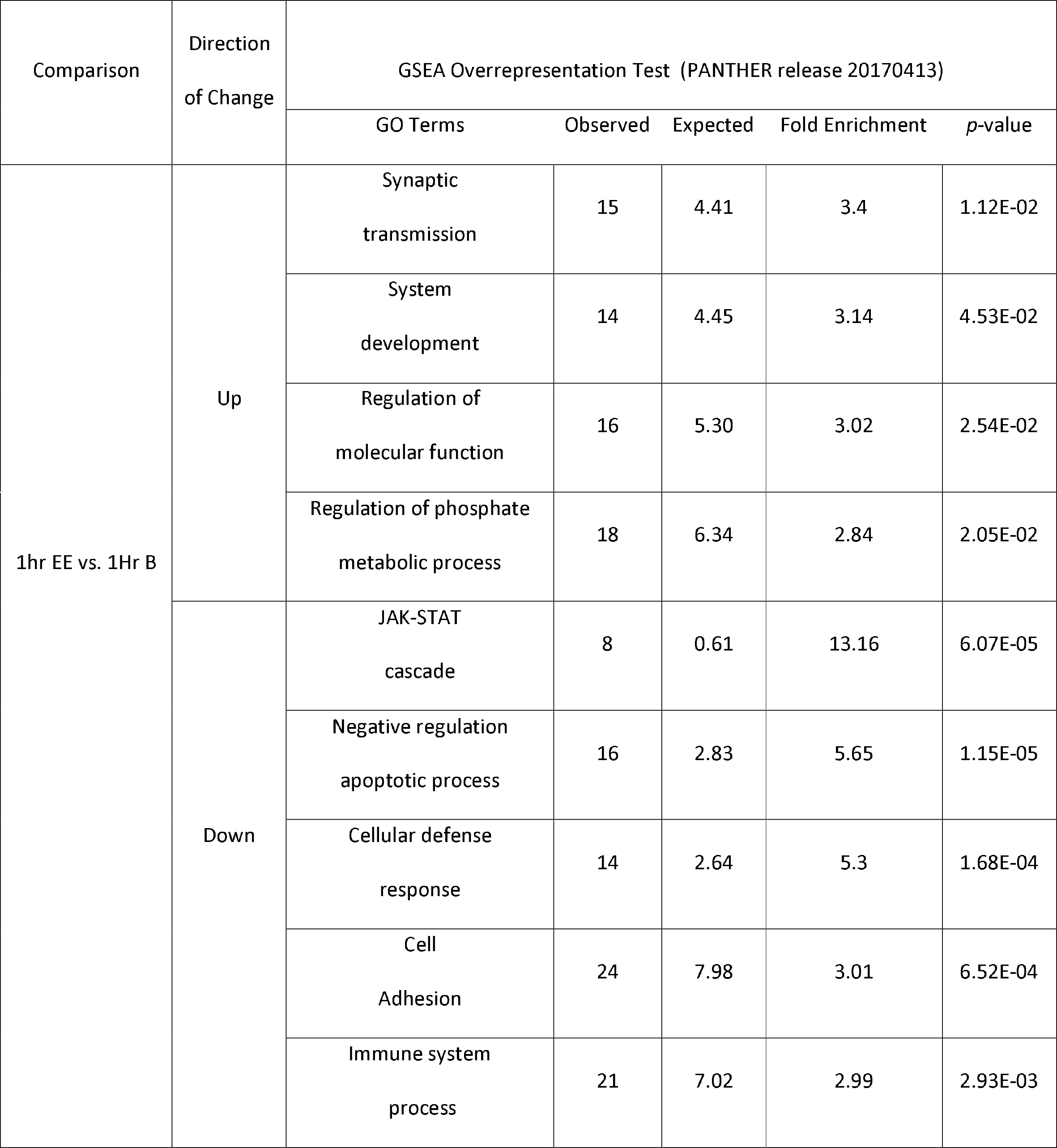
Panther GO-Slim Biological Process GO terms statistically overrepresented in the one hour EE vs one hour B comparisons. The ‘direction of change’ column signifies the direction of differential gene expression in the EE animals relative to the B animals. The observed column is the number of genes assigned this GO term in the submitted gene list; the expected column is the number of genes expected given the full *Sus scrofa* reference list.

Significant *p*-values from randomisation tests (up-regulated IEG signature *p*=9.99E-05, down-regulated microglial signature p=9.99E-05) would suggest these signatures do not occur by chance in EE animals.

In a network analysis using only the differentially expressed genes, the EE samples were close together in a relatively tight group, while the B samples were more dispersed, but still located together with little overlap with the EE samples (Figure 4). In addition there was no relationship with litter, indicating that for the differentially expressed gene set the effect of environment was stronger than any intrinsic difference based on genetics or early life (e.g. maternal) effects (Figure 5).

**Figure 5:**
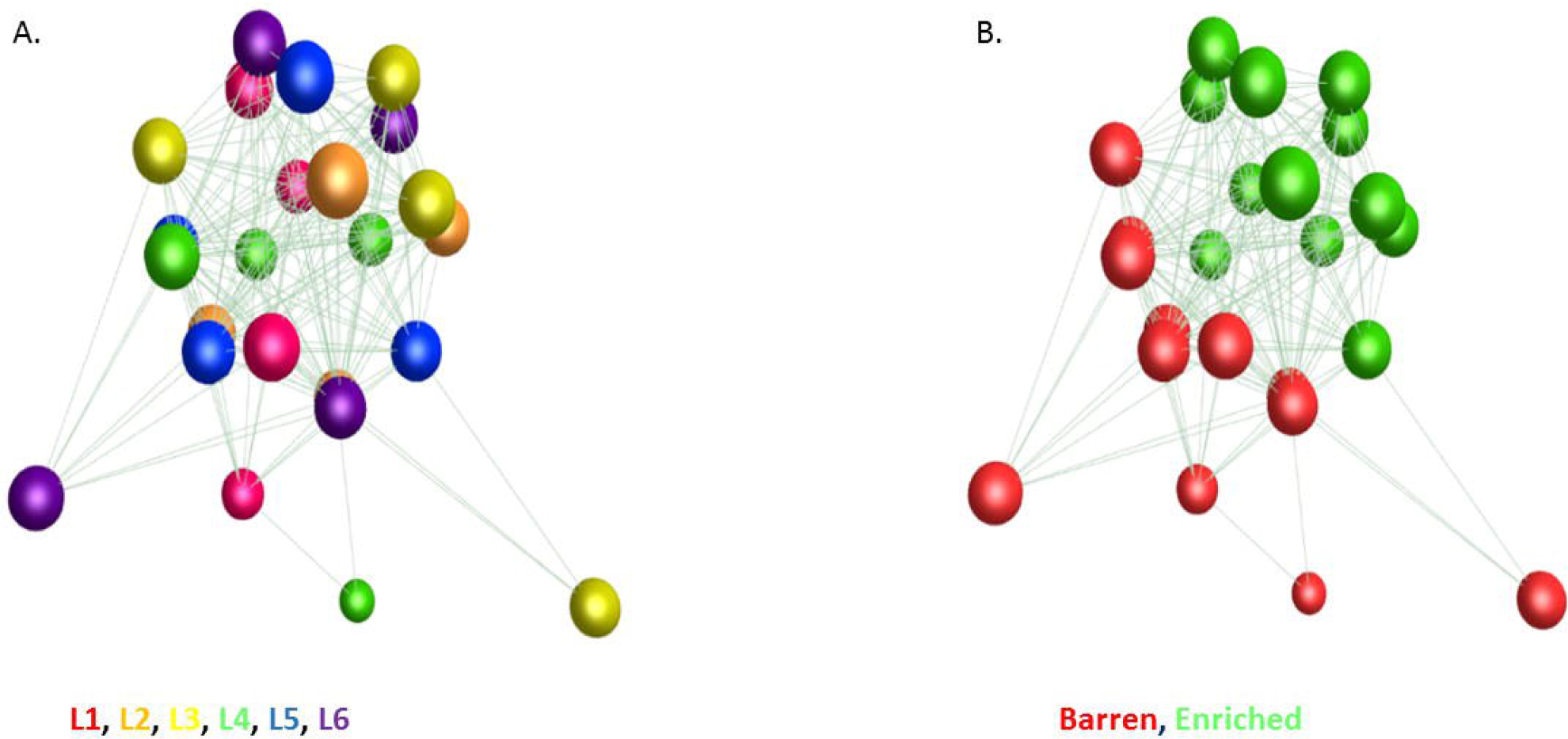
3D network graph of differentially expressed genes in the frontal cortex. Each node (sphere) in the network graph represents a sample, and each edge (line) a Pearson’s correlation between samples ≥ 0.93. There is no clustering of gene expression profiles based on litter of origin (Panel A) and an unambiguous effect of treatment on gene expression (Panel B). Inter-individual variability in expression profiles appears lower in the EE pigs than in the B pigs.

The expression profiles of the differentially expressed genes were analysed to create a gene-to-gene network. As might be expected (as these were differentially expressed in both directions) there were two clear elements in the network, comprising one element where the overall gene expression level was higher in the EE animals and a second element where the level was lower in EE animals. Similar results were seen when using the one hour, four hour or complete set of differentially expressed genes (Figure S1: Panel A-C segregated gene cluster histograms by animal, Panel D Clusters for all animals at 1 hour).

When the DEGs at one hour were used to construct a network graph, cluster 1 (which contains the majority of genes with average expression reduced in the EE condition) contained microglia-associated genes such as CSF1R (63), IRF8 (64) and P2RY12 (65) alongside various connective tissue genes including several collagen genes and FBN1. Cluster 4, where the average expression was increased in EE samples, contained IEGs such as ARC, several EGR genes, FOS and IER2. Cluster 2 contained largely neuronal genes which were not differently expressed between environments. Cluster 3 contained additional connective tissue genes with high levels in four of the B individuals. Clusters 5 and 6 represent groups of genes that were increased in a single individual. The gene lists for each cluster and average expression profiles for clusters where treatment differences were shown are presented in Table S3 and Figure S1.

The profiles for the four hour samples were largely determined by high or low expression in a single or small number of individuals. Many of the genes in this list were non-coding RNAs or unannotated genes. When both time points were included in the analysis, cluster 3 primarily contained IEGs and showed a pattern in which the one hour EE samples had higher expression than the one hour B samples, but at four hours the two groups had similar expression. One individual in the B group had a spike of IEG expression at four hours although most of the four hour B samples showed lower IEG expression than the EE samples.

## 4. Discussion

The EE piglets in this study displayed more active behaviour (combining locomotion (without object or social), social and object interaction) than B piglets, but only in the first hour after the additional enrichment stimulus (bag filled with straw) was provided. This was not affected by sex. This suggests that the piglets’ behavioural response to enrichment in this experiment was largely defined by the period that the additional enrichment stimulus remained novel. After the 4 hours of observation the piglets had tended to remove some or all of the straw from the enrichment stimulus and reduced the physical integrity of the bag which may have reduced the novelty value of the stimulus. Despite the larger space allowance and fresh bedding (straw, given as part of the additional enrichment stimulus) provided daily to the EE piglets, their overall activity profile was not different to the B housed piglets outside of the first hour post additional enrichment (using a scan sampling methodology). Overall total activity levels were lower in this study than in previous studies (see (66) for a review), but the proportion of active time spent interacting with the enrichment stimulus or the pen environment was higher than previously reported. Within the time spent active there were some effects of housing on specific behaviours. B housed piglets displayed more nosing of pen-mates (primarily nosing of the belly of other pigs) over the course of the observation, not dependent on time. This maladaptive behaviour is known to occur more frequently in piglets when the opportunities to use the snout to forage and explore are not provided(67). Despite no differences in time spent feeding being observed between treatments, piglets in the EE environment experienced higher growth rates than those in the B environment. Enriched housing has previously been shown to increase body weight and growth rate in grower pigs but not in piglets at such a young age (68). A recent meta-analysis has found that the provision of straw bedding is a significant factor in the increased average daily gain observed in EE housed pigs (69) though there has been little research to determine why this may be the case. In rodents, increased brain weight in the absence of increased body growth has been observed in environmentally enriched conditions (reviewed in (70)). Given the short time frame of this study we would not expect to observe significant differences in brain weight however over a longer period this may well be a contributory factor to observed weight gain.

The gene expression changes observed in this study mirror those of the behavioural differences, with most genes differentially expressed at one hour post the provision of the additional enrichment stimulation and few genes differentially expressed at four hours post his point. The observed increase in genes involved in synaptic transmission at the one hour time point in EE animals supports previous work describing a transient increase in cell activity in the hippocampus of rats following environmental enrichment (reviewed in (71)). This increase in synaptic transmission is likely to be the mechanism by which EE leads to increased brain weight (72), greater cortical depth (73), increased neurogenesis (6, 74) and increased synapse formation (75). The set of transiently-induced genes at one hour shows significant overlap with recent curated sets of IEGs defined in stimulated neurons based upon single cell RNA-seq (76, 77). Increases in expression of neurotrophic factor proteins (neuronal growth factors) have previously been observed in multiple brain regions of rats housed in enriched cages (14) and were observed as gene expression changes in the frontal cortex of the piglets in the current study, although not all growth factors fall within the gene set enrichment analysis. Neuronal growth factors (such as NGF, BDNF and VGF, all of which were upregulated in this sample set) are considered to be regulators of cell growth and survival in both the developing and adult nervous system (reviewed in(78)) and have been shown to acutely potentiate synaptic plasticity (79, 80).

Experience dependant plasticity, the way by which synaptic connections are strengthened or weakened, or indeed formed or eliminated, in response to experience is a well established mechanism (reviewed in(81)). Intracellular scaffold protein and regulator of membrane trafficking GRASP (mRNA upregulated in the EE pig frontal cortex) has multiple protein interacting domains allowing a wide range of protein assemblies to be trafficked at the synapse (82). GRASP is known to be involved in dendritic outgrowth (83) and in synaptic plasticity through its interaction with group I and II metabotrophic glutamate receptors (84). Reductions in GRASP protein expression have been observed in post mortem prefrontal cortex of schizophrenia patients (85) with no correlation to illness duration or history of medication. Similarly, decreases in mGluR2 have been observed in multiple psychiatric conditions including schizophrenia (86) and addiction (87), while treatment with the antidepressant medication imipramine has been shown to increase mGluR2 protein expression in the hippocampus of wild-type rats (88) but not to increase mRNA expression. Another G-protein coupled receptor upregulated in the EE pigs, GPR3, has been shown to be involved in anxiety related behaviours, with Gpr3-/- mice displaying increased anxiety-like behaviours that could be rescued with the use of the anxiolytic diazepam (89). Upregulation of these G-protein coupled receptors and their targets may suggest that EE piglets in the current study were experiencing reduced levels of anxiety and a more positive affective state compared to their B counterparts. Given the lack of sex difference in behavioural activity observed in this study, and no evidence of overlap between our list of DE genes and those shown to be sexually dimorphic in neonatal rodent cortex (90), we would propose that this effect would be consistent across both males and females.

An unexpectedly large proportion of the set of genes downregulated in the brains of the pigs exposed to environmental enrichment (relative to those in the B environment) are clearly associated with microglia. Numerous recent studies have identified microglia-associated genes in mice and humans (e.g. (59, 91–94)) however there has been little work focused on the pig. The set of genes down-regulated after one hour post the additional enrichment stimulation included generic myeloid markers such as PTPRC (CD45) and CD68, as well as most well-documented microglial markers (C1QA, MYD88, P2RY12, TGFBR2, TREM2). These findings suggest that microglia have altered transcriptional activity in EE relative to B pigs. With our study design it is impossible to determine whether the EE or B animals are the baseline so any changes in expression are relative, however irrespective of whether microglial activity is greater in B or reduced in EE animals, this result suggests that a relative reduction in activity may have benefits for brain health (95, 96). While reduced transcriptional activity could arise due to a selective reduction in microglia number, in rodents the number of microglia is relatively stable over time (97). Experimental depletion of microglia in rat neonates results in lifelong decreases in anxiety-like behaviours and increased locomotory behaviour (98), while increases in microglial numbers and activation states have been observed in stress-induced depression models (reviewed in (31)). In human patients, increased microglial activation has been observed using PET scanning in major depressive disorder (99), and from histology samples from brains of depressed suicides (100). This suggests that B housed animals may be experiencing greater anxiety and/or lower mood than their EE housed counterparts. This may provide part of the functional link as to why pigs housed in enriched environments display more optimistic judgement bias than those in barren conditions (101). Current evidence suggests that microglial gene expression is not sexually dimorphic at this equivalent developmental stage (102), and thus while only male brains were sampled in this study this may be extrapolated to include females with relative confidence.

Interestingly, at the four hour time-point there is a high proportion of 5s ribosomal and spliceosomal RNAs identified as differentially expressed (29% of the down regulated gene list, compared with 0.3% at one hour), which may be adding noise to the dataset in the later time-point. As observed in the network analysis, inter-individual variability in gene expression was greater at 4 hours, perhaps due to the varying rates of decay of the effect of the additional enrichment stimulus, or due to the animals no longer having a behavioural ‘focus’ (which may also explain the greater variability in gene expression between B individuals than between EE individuals). This is consistent with the behaviour analysis, which indicated that the benefit from the additional enrichment experience (bag of straw) was only felt while the object remained novel. This may contribute to the noise in the data at four hours, as animals may ‘cope’ with the lack of behavioural stimulation in different ways, or may be being stimulated by other factors within the home pen (such as other interactions with pen mates).

The current study would suggest the benefits of enrichment may occur in a pulse-like fashion, with bursts of behavioural activity being followed by a return to behavioural baseline, these bursts mirroring the active engagement with the behavioural opportunities offered by the additional enrichment stimulus. It would be of great interest to determine if these bursts of activity (and of related differential gene expression) alter the structure and function of the brain in a way that facilitates long term developmental change. Confirmation of the longer term effects of EE on microglial numbers and activation are required to determine if the observed gene expression changes are an indicator of lower neuronal health in B housed animals or if they are transient expression changes with little long term consequence.

## 5. Conclusions

In this study, behaviour was mainly organised around the time point when a daily enrichment stimulus was provided. This resulted in a ‘pulse’ of active behaviour that was mirrored by temporal changes in gene expression profiles in the frontal cortex. These gene expression changes primarily affected genes involved in synaptic plasticity, neuroprotection and the immune response, with a substantial fraction (15%) of known microglial signature genes showing relative down-regulation with enrichment. Our analyses suggest that relative to piglets in barren environments, those in enriched environments may experience reduced anxiety, increased neuroprotection and synaptic plasticity, and an immune response consistent with reduced inflammatory challenge.

## 6. Acknowledgements

We would like to acknowledge funding support for this work from the BBSRC Strategic funding to The Roslin Institute, and from the Scottish Government’s Rural and Environment Science and Analytical Services Division (RESAS). Technical support was provided by the SRUC technicians Marianne Farish Agnieszka Futro, Jo Donbavand and Mhairi Jack. Farm manager Peter Finnie provided general animal husbandry assistance throughout the experimental period. Behavioural data collection was performed by summer student Manon Poux and initial expression analysis by Edinburgh Genomics bioinformatician (Dr Frances Turner).

## 9. Supplemental Information

**Figure S1:** Cluster graphs from 1 hour Cluster lists found in Table S3 that show clear segregation between EE and B animals. Panel (A) Microglia and connective tissue genes; Panel (B) Immune and connective tissue genes; Panel (C) IEG; Panel (D) All clusters. Each column represents a single animal with B house pigs as columns 1-6 and EE pigs columns 7-12. There is clear segregation in panel D of the EE animals (top clusters) and B animals (bottom clusters)

**Table S1:** Means and SEM of behaviours at all time-points by treatment.

**Table S2:** Differentially expressed genes at 1 and 4 hours. With log2 fold change, p value, average counts and average transcripts per million for each treatment group.

**Table S3:** Genes present in gene-gene cluster analysis.

**Table S4:** IEG and microglial reference gene list. Reference list used for randomisation tests.

